# Parental care shapes the evolution of molecular genetic variation

**DOI:** 10.1101/2023.01.09.523216

**Authors:** R Mashoodh, A Trowsdale, A Manica, RM Kilner

**Affiliations:** Centre for Biodiversity & Environment Research, Department of Genetics, Evolution and Environment, University College London, United Kingdom; Department of Zoology, University of Cambridge, United Kingdom

**Keywords:** parental care, genetic variation, adaptation, mutation load, population genetics

## Abstract

Cooperative social behaviours, such as parental care, have long been hypothesized to relax selection leading to the accumulation of genetic variation in populations. Although the idea has been discussed for decades, there has been relatively little experimental work to investigate how social behaviour contributes to genetic variation in populations. Here, we investigate how parental care can shape molecular genetic variation in the subsocial insect *Nicrophorus vespilloides*. Using whole genome sequencing of populations that had evolved in the presence or absence of parental care for 30 generations, we show that parental care maintains levels of standing genetic variation. In contrast, under a harsh environment without care, strong directional selection caused a reduction in genetic variation. Furthermore, we show that adaptation to the loss of care is associated with genetic divergence between populations at loci related to stress, morphological development and transcriptional regulation. These data reveal how social behaviour is linked to the genetic processes that shape and maintain genetic diversity within populations, and provides rare empirical evidence for an old hypothesis.

**Lay Summary:** Social behaviours, such as parental care, have long been hypothesized to result in the accumulation of genetic variation in populations. Here, we investigate how parental care can shape molecular genetic variation in a species that perform biparental care, *Nicrophorus vespilloides*. Using genome sequencing of populations that had evolved in the presence or absence of parental care for 30 generations, we show that parental care maintains levels of standing genetic variation. In contrast, under a harsh environment without care, populations lost genetic variation. Furthermore, we show that adaptation to the loss of care is associated with genetic divergence between populations at genes related to stress, morphological development and transcriptional regulation. These data reveal how social behaviour is linked to the genetic processes that shape and maintain genetic diversity within populations.

## Introduction

While much recent work has focused on identifying genes that drive social behaviours [1,2], relatively few studies have examined the longstanding hypothesis that social behaviour affects the accumulation and maintenance of genetic variation. Yet, social living is associated with large-scale restructuring and the evolution of genome organization and architecture [3]. In humans, benevolent social activities, such as modern health care, are thought to have led to the accumulation of deleterious mutations within populations [4,5]. Therefore, the extent to which genetic variation is shaped by social behaviour has implications for the health of populations and their capacity to rapidly adapt to environmental perturbations. However, there have been few empirical tests of how social behaviour might drive genetic variation in practice. Here we investigate how a cooperative social behaviour, namely the supply of parental care, contributes to genome-wide levels of genetic variation. We focus on parental care in a subsocial pair-breeding insect, rather than more elaborate forms of sociality, to avoid the confounding effects of extreme reproductive skew on genetic variation, which is common in cooperative insect societies [6].

Cooperative social interactions often function to shield social partners from a harsh physical environment and the same is true for parental care [7,8]. Without cooperation generally, and care specifically, individuals would be exposed to strong, frequently directional, selection pressures from the abiotic environment, which would favour the evolution of new adaptations and cause an associated reduction in genetic variation. On the other hand, the presence of parental care relaxes selection from this wider environment, theoretically allowing genetic variation to accumulate. Indeed, several lines of evidence suggest that cooperative social behaviours, including care, can relax selection sufficiently to allow mildly deleterious mutations to accumulate within populations [9–13]. In this way, parental care, could shift the ‘mutation-selection’ balance by relaxing selection and preventing the elimination of new spontaneous mutations. The resulting increase in genetic variation could emerge in the form of single nucleotide polymorphisms (SNPs) and/or other structural genetic variants (e.g., indels, transposable elements and/or insertions/deletions) depending on the natural mutation rate of such variants. Exactly how care might maintain such variants has been the subject of some speculation [9]. One possibility is that “cryptic” variants could be maintained in the population with a combination of care-induced genetic capacitors, epigenetic modifications and/or RNA-mediated signals [14]. Nevertheless, although the suggestion that cooperative social behavour can shape genetic variation is relatively longstanding, we still have a poor understanding of how and where it might cause change at a molecular genetic level.

Here, we use evolving populations of burying beetles (*N. vespilloides*) to explore how parental care affects levels of standing genetic variation and how populations may adapt in the face of its loss. In natural populations of this locally abundant subsocial insect, burying beetle parents raise their young on a carrion nest, formed from a small dead animal, such as a mouse or songbird. There is continuous variation in the level of parental care supplied, with around 5% of parents abandoning the brood before their young have even hatched [15]. Offspring can survive without parental care, at least in the laboratory.

We exploited this natural variation in care to establish two types of experimentally evolving populations in the laboratory, which varied only in the family environment that larvae experienced during development, and where the same family environment was created for successive generations within populations. In Full Care populations (FC), parents remained with their young throughout development; whereas in No Care populations (NC), parents were removed just prior to hatching. No Care populations rapidly adapted to this regime (within 14 generations), with adaptive change being detectable through increases in breeding success and larval density (see Schrader et al., 2017). Moreover, we have previously shown that No Care populations evolved adaptively [16] and divergently from Full Care populations in the extent of the pre-hatching care behaviours [17], the extent of sibling cooperation [18,19] and in their larval morphology [20].

At the 30^th^ generation of experimental evolution, we use pooled whole-genome re-sequencing of these populations to document genetic variation at the molecular level when care was present and when it was prevented experimentally (**Figure 1a**). First, we determined the effect of care on within-population genetic variation (SNP diversity). Second, we identified the genetic loci that had diverged to the greatest extent following the removal of care by looking for regions of high genetic differentiation (F_ST_) between experimental populations.

**Figure 1.**
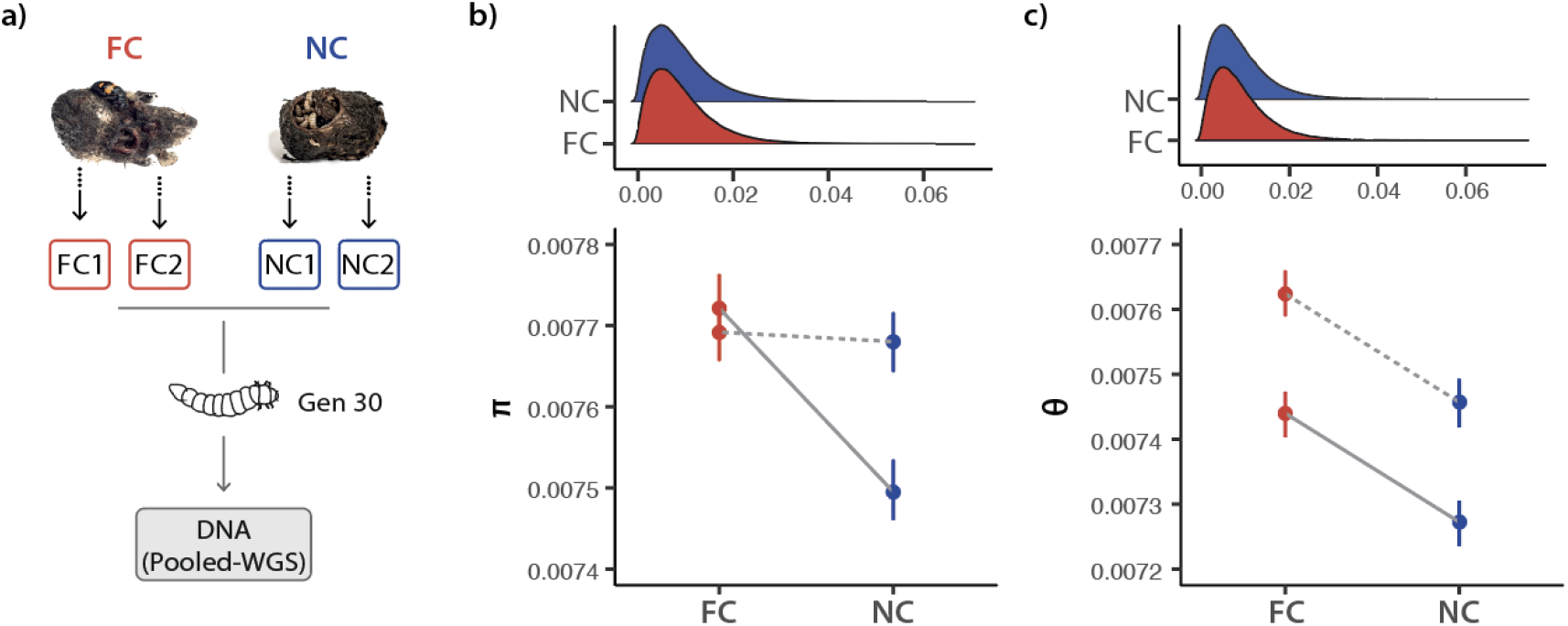
(a) Populations evolved in the presence (Full Care; FC) or in the absence of (No Care; NC) for 30 generations (two replicates per condition). Larvae were pooled for each replicate population (see Methods) for whole genome sequencing (WGS). Distribution (top) and median (bottom) of (b) Pi (π) and (c) Watterson’s theta (θ)across 1000bp non-overlapping windows for FC and NC populations (error bars represent 95% bootstrapped confidence intervals). Block 1 (dashed line) and Block 2 (solid line) are plotted separately.

## Methods

### Breeding design & Experimental Evolution

We sampled DNA from experimental populations of *Nicrophorus vespilloides* that had been evolving under different regimes of parental care, and which were founded from a single genetically diverse population generated by interbreeding beetles from multiple wild populations across Cambridgeshire. These populations have been described in detail previously [19], and comprise a total of 4 populations: two blocks (Block 1 and Block 2; separated by 1 week) containing two populations evolving with (FC_POP_) or without parental care (NC_POP_). For the first 14 generations, when directional selection was high, an average of 34 pairs of unrelated beetles were bred at each generation [19]. Thereafter, populations were maintained with an average of 37 and 49 pairs at each generation for FC and NC populations, respectively, and were equivalently successful across the generations (**Table S1**). On the 29^th^ generation (as in every generation previously), we paired sexually mature males and females within each population. Each pair was placed in a separate breeding box with moist soil and a thawed carcass (10-12g). We then placed each breeding box in a cupboard and allowed parents to prepare the carcass and for the female to lay the clutch of eggs. For the NC_POP_, after 53h, both parents were removed from the nest just as had occurred for the prior 29 generations. Approximately 80h after hatching we randomly selected 2-3 larvae from each family (15-18 families per population) for DNA extraction.

### Larval tissue dissection, DNA extraction & Whole-genome sequencing

For each family, DNA from first instar larvae were pooled and extracted using a modified version of the Qiagen DNEasy Mini Kit. Total DNA quality was checked using gel electrophoresis and yield was quantified using a Qubit DNA Assay Kit (Thermo Fisher). Families were pooled in equimolar concentrations such that each individual was represented equally to generate 4 libraries: FC1, FC2, NC1, and NC2 with pool sizes of 41, 52, 52, 59, respectively. Whole genome re-sequencing libraries were constructed and sequenced (150bp paired-end) at a depth of 100x using an Illumina Novaseq 6000 platform by Novogene (Hong Kong).

### Bioinformatic Analyses

Reads were trimmed using TrimGalore (0.5.0; https://github.com/FelixKrueger/TrimGalore) to remove adaptor sequences, perform quality trimming and discard low-quality reads. Reads were aligned in paired-end mode using the burrows-wheeler aligner (bwa) to the *N. vespilloides* reference genome (NCBI Refseq Assembly: GCF_001412225.1) [21,22]. See **Table S2** for read mapping statistics. Duplicates were removed using PicardTools (http://broadinstitute.github.io/picard/). Pileup files were created using *samtools* [23] from mapped reads and indels and repeats were filtered using the *Popoolation* toolbox [24]. These pileup files were used to calculate measures of genetic diversity with *Popoolation* (π, Watterson’s θ, synonymous vs non-synonymous rate of π, and Tajima’s D). Pileups were were merged into a single sync file using *Popoolation2* for use with *poolfstat* [25], *Baypass* [26] and *Popoolation2* to measure between-population divergence (e.g., F_ST_, Fishers’ Exact Tests and Bayesian auxillary models; described below). All subsequent post-processing and statistical analyses were performed in *R* version 4.1.2 using the core *R stats* package [27]. Data wrangling and visualisations were performed using the *tidyverse* suite [28].

#### Intrapopulation genetic variation

We used π and Watterson’s θ to measure levels of standing genetic variation within populations. Watterson’s θ represents the expected number of segregating sites observed between a pair of homologous sequences sampled from a given population, whereas π is the average number of pairwise difference between all possible pairs of individuals in the sample. These measures were calculated for non-overlapping 1000bp windows (for sites with coverage between 40-700 reads) across the genome using tools from *Popoolation*. We also computed gene-wise synonymous and non-synonymous pi for CDS coordinates of all genes extracted from the reference annotation using *Popoolation*. To allow comparisons to F_ST_ windows we computed Tajima’s D for 500bp sliding windows with a 250bp overlap. For all genetic diversity measures, we used non-parametric Kruskal-Wallis tests to test for differences between Full Care and No Care populations separately for each replicate block except in the case of Tajima’s D. For normally-distributed Tajima’s D values we used t-tests to test for differences between care conditions within each block. Windows were filtered so that statistics were based on windows that were covered across all replicates of both populations.

#### Genetic divergence between populations

To estimate population structure and demographic history, we extracted SNPs from the population sync file using the R package *poolfstat* [25] using the core model of *BayPass* version 2.3 [29]. Baypass uses allele-frequencies to estimate a scaled covariance (Ω) matrix, which can be interpreted as the pairwise estimates of differentiation between the population. The Ω matrix was converted to a corrrelation matrix in R and visusalised as a tree using the base R *stats* package.

To further measure the extent of genetic divergence between populations, we used *Popoolation2* [30] to calculate the pairwise fixation index (F_ST_) for all combinations of population pairs across 500bp sliding windows (250bp overlap) across the genome. SNPs were called using sites with read counts between 40 and 700. Hierarchical clustering indicated that NC1 and NC2 were more closely related to their FC counterparts than to each other (**Figure S1**). Moreover, inspection of F_ST_ values across the genome indicated that the overall magnitude of differences between FC and NC differed between the blocks (**Figure S2**). Therefore, to identify windows where evolving populations may have diverged consistently, over and above any variation within and between blocks, we computed F_ST_ for each replicate line separately (i.e., FC1;NC1 and FC2;NC2). We then performed Fisher’s Exact Tests for each of these windows to screen for significant allele frequency differences. We took the product of the -log(*p*) values for each block (FC1;NC1 *x* FC2;NC2) and selected the top 0.5% of values as regions of interest. In this way, we selected for loci which diverged consistently across the blocks, assuming that inconsistent divergence may reflect drift. Location of windows of interest were annotated using the reference genome and the *intersect* command in *bedtools* [31]. A hit was considered only if the window intersected with the coordinates of the annotation by at least 1bp.

Using the same logic we used the auxillary model in BayPass to identify candidate SNPs that were consistently associated with the loss of care across both blocks. Using the covariance structure among the population allele frequencies (Ω), the model explicitly accounts for the shared history of the populations, rendering the identification of SNPs potentially subjected to selection less sensitive to the confounding effect of demography [26,32]. Specifically, the model involves the introduction of a binary auxiliary variable to classify each locus as being associated or not with the loss of care. This allows the estimation of posterior inclusion probabilities (and Bayes factors) for each SNP while also accounting for multiple testing issues. For each SNP, the Bayes factor was expressed in deciban units (dB) via the transformation 10log10(BF). Significance was assessed based on the Bayes Factor (BF) between models and SNP markers with strong evidence (BF > 20) were retained as potential candidates of interest (according to Jeffrey’s rule) [33]. We then examined where these SNPs were located by looking for genes within 500bp of the outlier SNP (using *bedtools* ‘window’) making this comparable to our windowed approach.

### Functional Annotation

Functional enrichment analyses were conducted using the topGO R package version 2.38.1 [34] to identify over-representation of particular functional groups within the diverged genes in response to the removal of care, based on GO classifications using Fisher’s exact test. GO terms were annotated to the *N. vespilloides* genome using the BLAST2GO (version 5.1.1) workflow to assign homologs to the *Drosophila* non-redundant protein databases [35]. To improve the GO term assignment, *N. vespilloides* genes were further annotated using a custom script that assigned GO terms from multiple well-annotated insect species (e.g., *A. mellifera, B. terrestris, A. cephalotes, N. vitripennis, T. castaneum* and *O. taurus*) based on ortholog assignments obtained using Orthofinder [36] using a custom pipeline (https://github.com/chriswyatt1/Goatee). To identify transcription factors we searched for the presence of known Pfam [37] transcription factor domains in the protein sequences of the gene candidates of interest using Interproscan [38]. Putative promoter regions (5’ UTRs) were classified as the 500bp region upstream of each gene start coordinate [31].

## Results

### Standing genetic variation between populations

First, we determined the effect of care on within-population standing genetic variation by measuring genetic diversity. We computed both Watterson’s theta (θ) and Pi (π) statistics for each population across 1000bp non-overlapping windows. Populations that evolved under Full Care (FC1 and FC2) had higher theta values than populations evolved under No Care (NC1 and NC2) (**Table 1**; Figure **1c**; all *p*’s < 0.001). Similarly, there were higher Pi values in FC compared to NC, though this effect was not present in Block 1 (**Table 1**; **Figure 1b**). Together, these results suggest that FC populations maintained more SNP diversity compared to populations evolving under NC with some detectable variation between blocks **(Table 1**). We measured Tajima’s D (500 bp overlapping windows) to further characterize the evolutionary forces shaping genetic diversity between populations. We show that genome-wide levels of Tajima’s D are negative, with both replicates showing a significant reduction in Tajima’s D in NC compared to FC (**Table 2**; **Figure 3**).

**Table 1.**
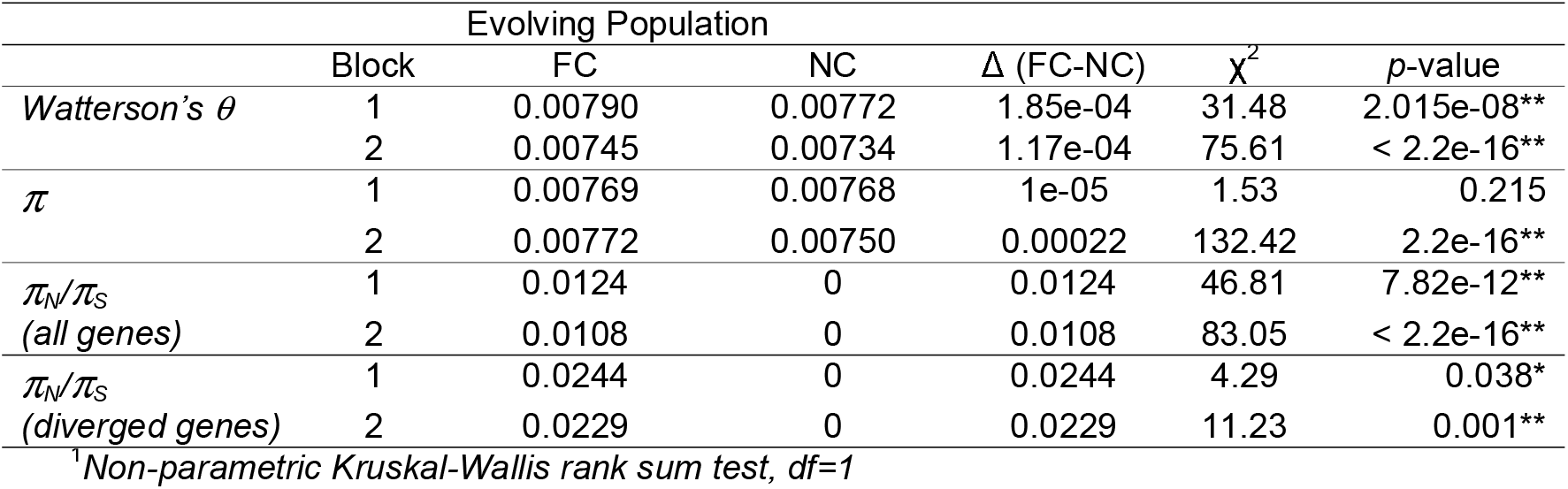
Genetic diversity measures for each population evolving under Full Care (FC) and No Care (NC) for each block. Delta is the difference between FC and NC populations computed separately for each block (* indicates p < 0.05, ** indicates p< 0.001).

**Table 2.** Tajima’s D (mean of 500bp sliding windows) for populations evolving under Full Care (FC) and No Care (NC) for each block. Statistics are presented for all windows across the genome (genome-wide) as well as for windows that overlapped with diverged genes (+/-5kb). Delta is difference between Tajima’s D between means of FC and NC populations (** indicates p < 0.001).

|  | Evolving Population |  |  |  |  |  |
| --- | --- | --- | --- | --- | --- | --- |
| | Block | FC | NC | $\Delta$ (FC-NC) | t (df) | p-value |
| Tajima's D (genome-wide) | 1 | -0.134 | -0.155 | 0.020 | 11.63<sup>1</sup> | < 2.2e-16* |
|  | 2 | -0.007 | -0.053 | 0.046 | 26.16<sup>2</sup> | < 2.2e-16* |
| Tajima's D (diverged genes) | 1 | 0.039 | 0.024 | 0.086 | 3.22<sup>3</sup> | 0.001* |
|  | 2 | 0.205 | 0.119 | 0.015 | 17.59<sup>4</sup> | < 2.2e-16* |

<sup>1</sup>df=1288045 <sup>2</sup>df=1288388 <sup>3</sup>df=167272 <sup>4</sup>df=167250

### Genetic differences between populations

Next, we identified the genetic loci that had diverged to the greatest extent following the removal of care by looking for regions of high genetic differentiation (F_ST_) between experimental populations (see **Figure S1** for population structure). We looked for changes over and above drift by looking for highly consistent divergence across the replicates using a 500bp sliding window approach (see Methods; **Figure 2a**). Highly significant windows overlapped with both protein-coding and regulatory features of the genome, with 16% of windows being classified as a regulatory change in contrast to 47.9% of windows in protein coding genes (**Figure 2b**). Using this approach, we identified 648 differentiated genes (**Table S3**), with 144 of these windows uniquely intersecting the 5’ UTR regions of these genes only (**Figure 2b**; **Table S4)**. These genes were generally enriched for GO processes associated with morphogenesis, neural development, immunity, and hormone signaling (**Figure 2c**; **Table S5**). To test that our approach converged with other methods, we also identified SNP outliers using a Bayesian approach (see Methods). This method identified 3086 outlier SNPs with consistent allele frequency differences between the NC and FC populations across both replicate blocks, which fell within 500bp of 1176 genes (**Table S6**). These SNP outliers broadly converged on our windowed approach (220 genes; **Figure S3**) with several key genes identified in both methods (**Table S7**).

**Figure 2.**
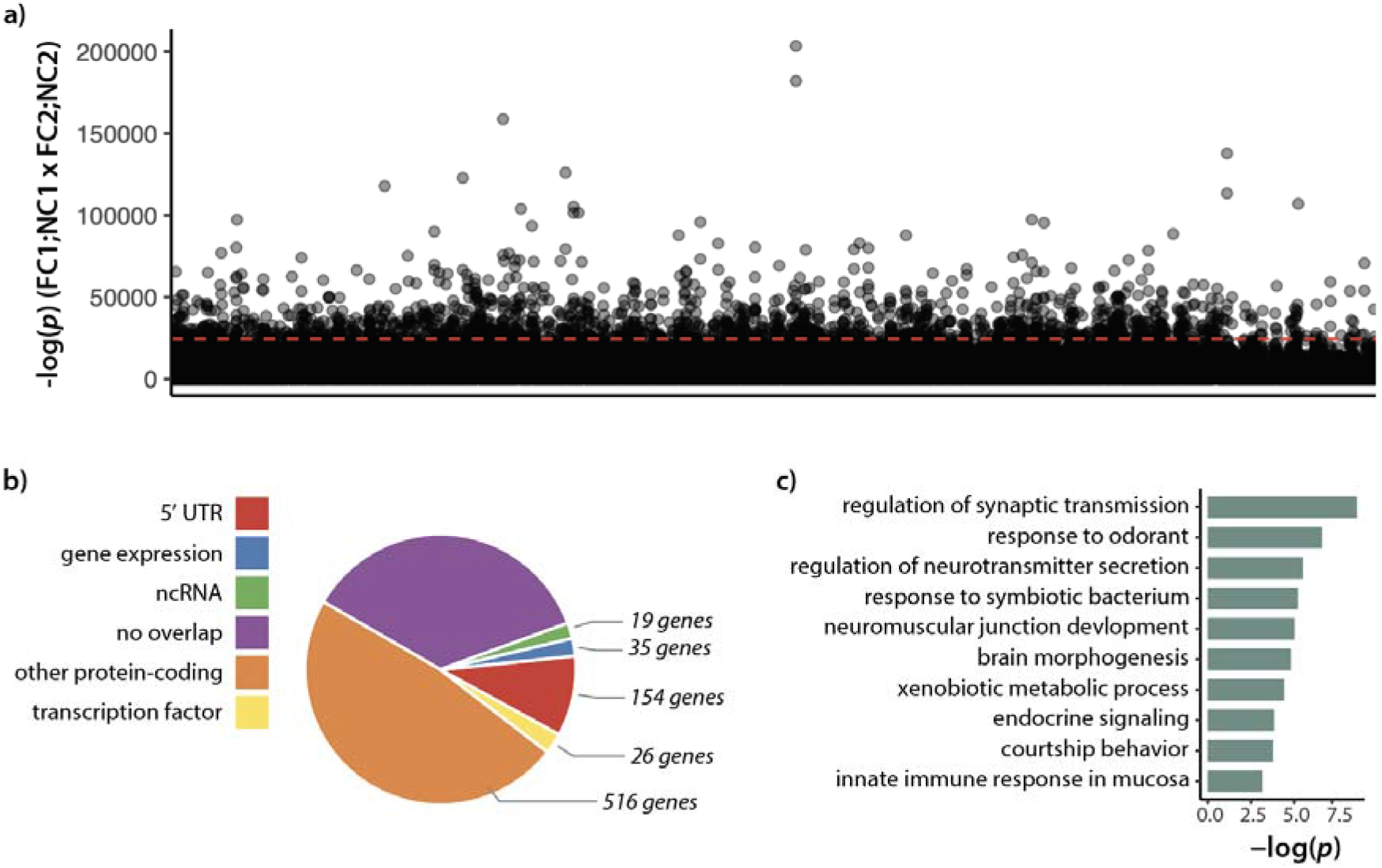
(a) Product of -log(*p*) values for allele frequency differences between Full Care (FC) and No Care (NC) populations of each block (i.e., FC1;NC1 *x* FC2;NC2) for 500bp sliding windows (250bp overlap) sorted by position. Dashed red line indicates 99.5^th^ percentile (b) Percent of windows overlapping with genomic features (ncRNA = non-coding RNA; 5’ UTR is defined as 500bp upstream of gene start position) and the number of genes that correspond to each category (c) Representative enriched GO terms (biological processes) for the most diverged genes between FC and NC populations.

To test the hypothesis that genes selected in the No Care lost genetic variation, we examined the gene-wise ratio of non-synonymous to synonymous π (π_N_/π_S_; **Table 1 & Figure S4**) and Tajima’s D within 5kb of divergent loci (**Table 2; Figure 3**). Both measures were reduced in No Care populations relative to the Full Care populations, in both replicate blocks, and this was true genome-wide as well as for the genes identified in our divergence screens.

**Figure 3.**
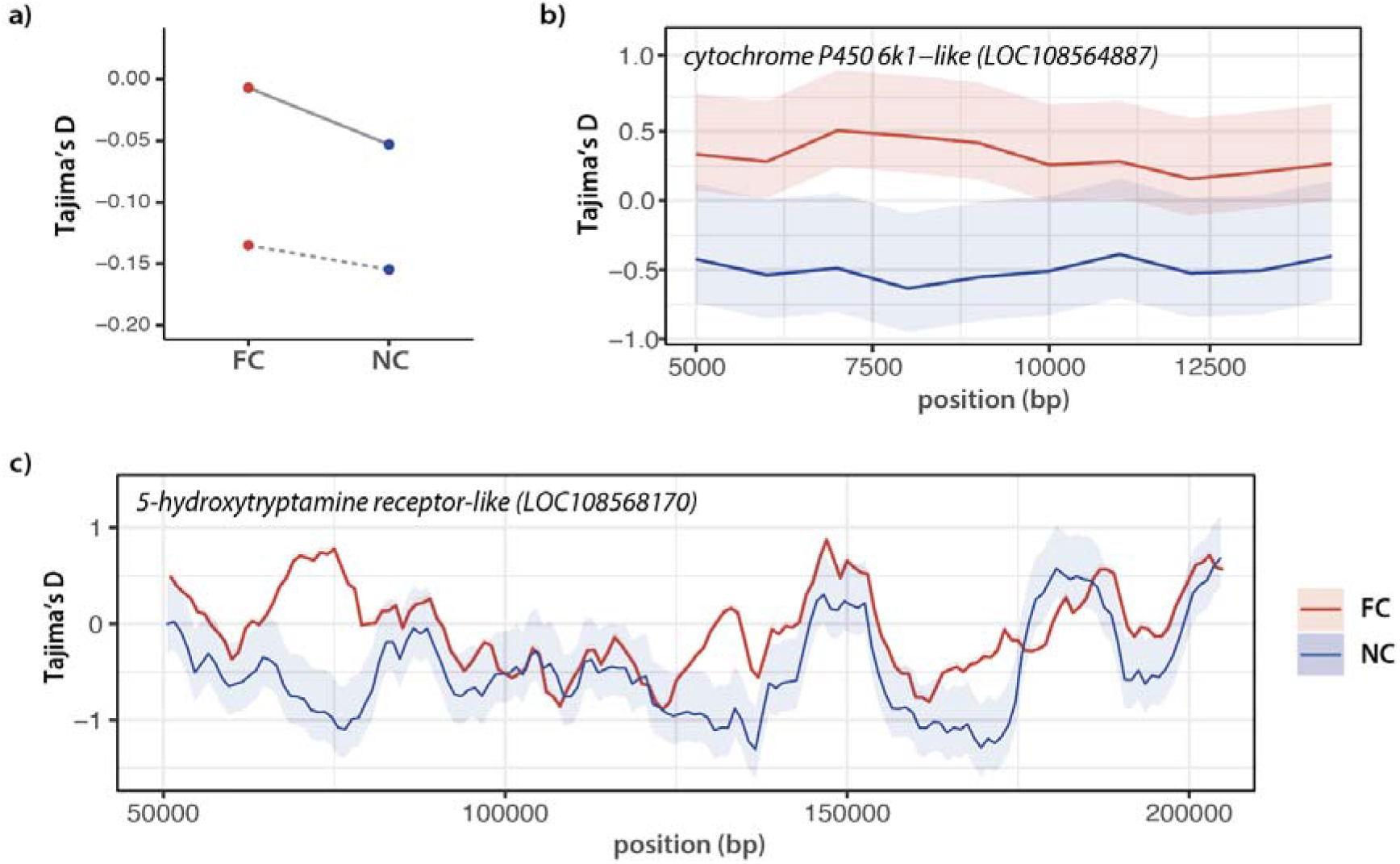
(a) Genome-wide Tajima’s D (mean) for 500bp sliding windows for No Care (NC) and Full Care (FC) populations (Block 1: dashed; Block 2: solid). Average Tajima’s D for FC and NC populations along gene bodies of two genes that showed extreme Tajima’s D values (bottom 5%) and showed allele frequency divergence (Table S3) (b) cytochrome P450 6k1-like and (c) 5-hydroxytryptamine receptor-like. See Figure S5 for replicate blocks plotted separately. All error bars represent 95% bootstrapped confidence intervals.

## Discussion

We found that populations with parental care (Full Care) had greater levels of genetic variation, in the form of higher theta and pi diversity, than the populations where care was prevented (No Care). Previous work has suggested that social behaviour contributes to genetic diversity mainly because of its effect on demography, and particularly because of its influence on effective population size (Ne). Population genetic theory predicts that genetic diversity will increase with Ne and mutation rate [39]. Previous empirical work linking behavioural and life-history traits, such as reproductive strategy, fecundity and body size with genetic diversity has suggested that these associations are ultimately mediated by changes in Ne [40–43]. However, our results cannot be explained by demography because populations were maintained at similar population sizes with no differences in fecundity [44] and no possibility of overlapping generations and/or changes in mating structure. Any small deviations in population size between care treatments were biased towards reducing genetic diversity in the Full Care populations – yet we found the opposite result. We suggest instead that the accumulation of genetic variation here is due directly to the effect of parental care in relaxing selection. The founding wild populations were inclined to provide care [20] and likely had already accumulated high levels of standing genetic variation, which was swiftly lost when we exposed populations to selection in a No Care environment.

The majority of this accumulated genetic variation is likely to be either neutral or mildly deleterious, since the majority of new mutations generally fall into either of these two categories [5,45]. Indeed, we have previously demonstrated that inbreeding of these populations resulted in faster extinction of Full Care compared to No Care populations, further suggesting that, at least some, of the variation accumulated in the presence of care was deleterious [12]. Although we measured only SNP variation here, genetic variants (e.g., insertions/deletions, transpositions) that arise through different types of mutation or recombination could also, in theory, be maintained in the population by parental care. Whether care favours particular types of mutants remains to be tested in future studies. In contrast, the harsher No Care environment imposed strong directional selection resulting in rapid adaptation [16,19] and reducing levels of standing genetic variation. We identified genetic divergence at number of loci, which were also associated with the loss of non-synonymous pi (lower π_N_/π_S_) and reductions in Tajima’s D, a pattern that was similar to the genome-wide differences in genetic diversity. Again, this is consistent with the interpretation that No Care populations experienced strong directional selection, whilst Full Care populations harboured more potentially deleterious mutations.

Here, we follow convention in assuming that loci that diverge consistently are likely to represent adaptive genomic change, whereas inconsistent responses to the parental care treatment (No Care versus Full Care) are due to drift. Yet alternative explanations for inconsistent responses are also possible, and this might be particularly true for traits under social selection (as opposed to abiotic selection pressures). Inconsistent patterns of genetic change across experimental blocks attributed to drift might instead reflect idiosyncratic or opportunistic responses to selection that arise through subtle variation in founding populations [46–48]. This is not surprising given that polygenic traits can be genetically redundant and adaptation can arise through multiple intersecting pathways and unique combinations of alleles within a population [46,49]. Moreover, the magnitude of these inconsistent differences, either due to drift or selection, might have been intensified by the selection regime imposed by the social environment, depending on whether it relaxed selection or imposed directional selection, for example, or whether the strength of selection was modulated by genes of social partners, parents or siblings [10,50]. Such effects could explain variation within and between replicate populations that accumulates over time. Although we cannot distinguish idiosyncratic adaptive change from drift with our data currently, future work using a high number of replicated populations measured across several generations could provide key insights into these evolutionary dynamics [46].

Our data suggest that No Care populations diverge from Full Care populations at loci that could promote immunity, metabolic and behavioural stress resilience in the absence of care. The loss of care in *N. vespilloides* is likely to be associated with greater levels of environmental stress during development and heightened exposure to pathogens from the carrion resource [51,52]. We have previously shown that adaptation to a No Care environment is associated with gene expression signatures that show blunted stress responses and compensatory expression in metabolic and developmental pathways [51]. Not surprisingly, many of the genetic differences between the populations are in upstream regions and/or genes that encode for transcription factors or chromatin modifiers, suggesting that change in regulatory function is a key component of adaptation to the loss of care. This is likely to be an underestimate of the extent of regulatory change, as we have yet to characterise the regulatory landscape of *N. vespilloides* and windows without an annotated overlap could be in distal promoter and/or enhancer regions. Nevertheless, differences in regulatory functions could shape levels of gene expression of other genes, further buffering against stress in the absence of parental care [51]. In this way, the signatures of adaptation to the loss of parental care are not much different to adaptive genetic responses to other abiotic stressors in the broader environment. Indeed, a key feature of stress adaptation across species is that it involves changes in gene regulatory pathways and this is true from bacteria to plants and animals [46,53–55]. Although we cannot identify a single gene or master regulator within the regulatory changes, these data do identify candidate regulatory genes that might play key roles in conferring resilience to the loss of care, and to environmental stressors more broadly.

Delving more deeply into loci at which we detected the greatest differences, we found that adaptation to the loss of care involved changes in several genes associated with immune function (e.g., *CD109 antigen* and *lysozyme c-1*), which could help cope with the increased exposure to the bacterial pathogens of the carcass nest experienced by No Care larvae. Previous work on *N. vespilloides* has shown that lysozyme expression is particularly heightened in parents immediately after the larvae hatch [56,57] and that it is likely to be particularly important for eliminating pathogenic Gammaproteobacteria [58]. Relatedly, we found divergence at a number of cytochrome P450 genes (*4ac1, 4ac2, 4c1, 4g15, 9e3 and 6k1-like*; Table S3), which are known to be involved in the metabolism of endogenous compounds as well as exogenous toxins and disease vectors, and which might also participate in defensive responses [59].

Cytochrome genes also appear to play a role in mediating the response to social density in Drosophila. Expression changes at the *Cyp4, Cyp6* and *Cyp9* gene families can be induced by manipulating social density in *Drosophila* and deletions of the *Cyp6a20* gene have been associated with higher levels of aggression and reduced sociality [60]. This is particularly interesting given that we have previously shown that larvae from the No Care populations evolved to show greater levels of sibling cooperation than larvae from the Full Care populations [13,18]. Cytochrome P450 families tend to share similar functional domains, and therefore, it is possible that changes in these genes could have effects on social behaviour *via* its actions on multiple hormonal systems (e.g., pheromones, ecdysone) [59,61]. Furthermore, P450 genes are also intertwined with juvenile hormone pathways which are known to be involved in multiple facets of behavioural and morphological development [62]. This could include adaptations that aid in locating and facilitating the use of the carrion breeding resource, such as the increase in relative mandible size and reduced arrival time at the carcass that we also detected in No Care larvae [20,63]. This interpretation is additionally consistent with most of the genetic hits belonging to cell signaling and biosynthetic pathways which fall into GO categories associated with morphological, brain and olfactory development.

We also found changes in neuropeptides that are also involved in metabolic, homeostatic and feeding pathways (e.g., *orexin, 5-hydroxytryptamine and cholecystokinin receptors*), raising the possibility that these could represent adaptations in larvae for feeding and extracting nutrients from the carcass resource in the absence of parents [64]. Previous work has shown that genes ancestrally associated with metabolic, homeostatic and feeding pathways can be co-opted to serve new social functions [65,66]. For example, the oxytocin/vasopressin system is commonly associated with the expression of parental care, pair bonding and other affiliative social behaviours in mammals [67], but has an ancestral role associated with promoting water balance [68]. A recent study in the closely related burying beetle, *N. orbicollis*, suggested that the expression of *inotocin* (the insect homologue of oxytocin/vasopressin) was correlated with the transition to parenting, an effect that was more pronounced in males than females [65]. *Takeout* is another gene that, despite being typically associated with feeding and circadian rhythms, has been shown to be highly expressed whilst parenting in burying beetles and may be involved in the transition from infanticide to larval care [66,69,70]. We found divergence in both an *oxytocin receptor* and a *takeout* homologue (Table S3), which could explain how male parental care eventually decayed in the No Care lines [71]. Finally, *angiotensin converting enzyme* was one of the most diverged genes in our analyses. This gene has varied roles from conferring immunity to regulating neuropeptide signaling [72]. However, it also appears to be strongly expressed in insect reproductive tissues and may, therefore, play a role in adaptations in mating and fecundity between the populations [44].

While we show here that parental care contributes to genetic variation through its effect on selection, it is possible that the incidence of mutation is itself reduced by the loss of parental care. Mutation rates have a strong genetic basis and can vary between individuals and amongst populations [73,74]. Given that the loss of care is a major developmental stressor, and that stress has been shown to induce mutations, adaptation to the loss of care could involve genetic mechanisms that dampen and/or buffer the consequences of new mutations that arise [9,45]. Consistent with this hypothesis, we found high levels of genetic differentiation amongst genes involved in DNA replication and repair (e.g., *Artemis* and the *PAXIP1 interacting protein*; Table S3) [75,76]. These genes, as part of their role in stress regulation, could facilitate efficient DNA repair, purging new mutations and shaping the subsequent mutation load of a population. The observation that multiple transfer RNAs (tRNAs) show divergence is particularly interesting given that variation in tRNAs have been associated with increased mutation loads *via* transcription assisted mutagenesis [77]. In other words, both genetic and phenotypic adaptations within each population could favour an optimal mutation-selection balance, resulting in different levels of standing genetic variation based on levels of care experienced within each population. Further functional characterization of these changes would help clarify if parental care facilitates the evolution of the mutation rate, potentially providing another mechanism for divergence in mutation-selection balance among populations [45,78].

In short, we have shown that parental care allows genetic variation to accumulate by relaxing selection. When care is lost, a number of genetic changes quickly follow which may be adaptive and which result in the loss of standing genetic variation. Better functional characterization of these gene targets and regulatory regions is now required to understand the genetic causes and functional consequences of the differences we have found between populations that are, and are not, exposed to post-hatching care. These are key areas of future work that will help explain whether the maintenance of standing genetic variation under parental care is likely to help or hinder adaptation in a rapidly changing world.

## Data Access & Code Availability

All raw sequencing data generated have been submitted to the Sequence Read Archive (SRA) under BioProject ID: PRJNA934336. All code for the analyses contained within this manuscript can be found at: https://github.com/r-mashoodh/nves_dnaEvol

## Acknowledgements

We thank Benjamin Jarrett and Darren Rebar for their help with beetle population maintenance. The molecular aspects of this project were supported by a Biotechnology and Biological Sciences Research Council Future Leaders Fellowship (BB/R01115X/1) and a Royal Society Small Research Grant (RGS\R1\191162) to RM. The experimental population work was supported by a Consolidator’s Grant from the European Research Council (310785 Baldwinian_Beetles), by a Wolfson Merit Award from the Royal Society and The Leverhulme Trust (RPG-2018-232) each to RMK. This work was also supported by a Newton Trust grant (20.40(b)) held jointly by RM and RMK. Finally, we would like to thank the editors, Allen Moore and two anonymous reviewers for their thoughtful comments which significantly improved the manuscript.

## Author Contributions

RM and RMK conceived of the study. RM and AT conducted the experimental work. RM and AM analysed the data. RM wrote the paper. All authors discussed the results and commented on the manuscript.

