## Supplementary Information for "Parental care shapes the evolution of molecular genetic variation"

**for**

*Corresponding author

**Table S1.** The number of pairs bred (number successful) for No Care (NC1 and NC2) and Full Care (FC1 and FC2) populations from generation 15 until the sampling of DNA for the current study. Data for generations 1 to 14 are presented in Schrader et al., 2017 [19].

| Generation | FC1 | FC2 | NC1 | NC2 |
| --- | --- | --- | --- | --- |
| 15 | 35 (35) | 35 (34) | 50 (36) | 50 (43) |
| 16 | 35 (31) | 35 (19) | 50 (46) | 50 (37) |
| 17 | 50 (46) | 49 (44) | 50 (44) | 50 (38) |
| 18 | 35 (31) | 35 (30) | 35 (31) | 35 (35) |
| 19 | 35 (31) | 35 (31) | 50 (35) | 50 (44) |
| 20 | 35 (27) | 35 (33) | 50 (30) | 50 (38) |
| 21 | 49 (47) | 49 (46) | 50 (37) | 50 (46) |
| 22 | 35 (34) | 35 (33) | 50 (38) | 50 (36) |
| 23 | 35 (33) | 35 (35) | 50 (36) | 50 (31) |
| 24 | 40 (37) | 40 (38) | 45 (43) | 45 (45) |
| 25 | 35 (33) | 35 (31) | 50 (39) | 50 (43) |
| 26 | 35 (34) | 35 (32) | 50 (48) | 50 (42) |
| 27 | 35 (34) | 35 (31) | 50 (26) | 50 (31) |
| 28 | 35 (25) | 35 (21) | 60 (32) | 60 (44) |
| 29 | 40 (34) | 44 (43) | 45 (42) | 44 (41) |
| 30 | 35 (31) | 35 (31) | 50 (43) | 50 (45) |
| mean pairs | 37.4 | 37.6 | 49.1 | 49 |
| mean survival (%) | 90 | 88 | 78 | 82 |

**Table S2.** Mapping statistics of pooled libraries for both replicates of populations evolving in the presence of parental care (Full Care; FC1 and FC2) and in the absence of care (No Care, NC1 and NC2).

| Population | Total reads (count) | Mapped reads (count) | Mapping rate (%) | Average depth (X) |
| --- | --- | --- | --- | --- |
| FC1 | 140,310,148 | 122,522,219 | 87.32 | 79.86 |
| FC2 | 163,923,157 | 143,606,777 | 87.61 | 93.34 |
| NC1 | 163,237,789 | 140,227,292 | 85.90 | 91.55 |
| NC2 | 151,459,484 | 131,781,027 | 87.01 | 85.11 |

*The following tables are included as attachments:*

**Table S3.** Top 0.5% of genes showing consistent divergence across the populations.

**Table S4.** List of 5’ UTRs in outlier windows.

**Table S5.** List of significant GO terms for genes that are divergent between populations.

**Table S6**. Outlier SNPs, allele frequencies and closest gene within 500bp.

**Table S7**. Overlapping genes between window outlier and SNP outlier approach

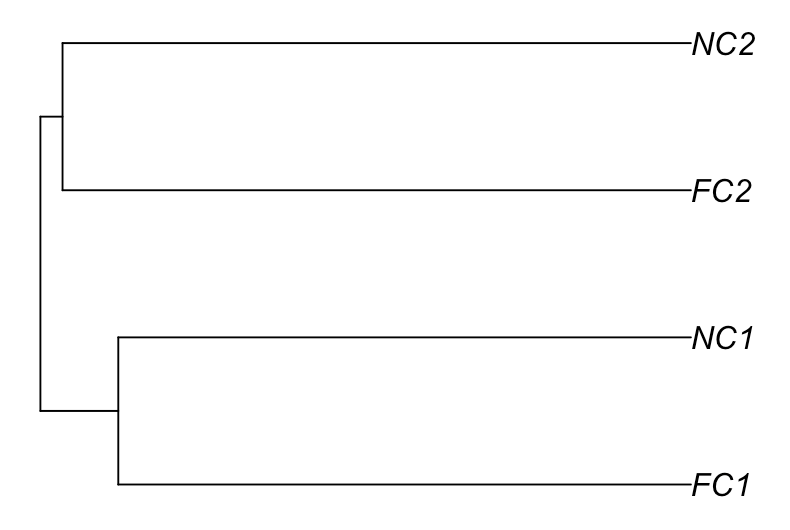

**Figure S1.** Population structure, based on hierarchical clustering of a scaled SNP covariance matrix derived for populations for both replicates of populations evolving in the presence of parental care (Full Care; FC1 and FC2) and in the absence of care (No Care, NC1 and NC2).

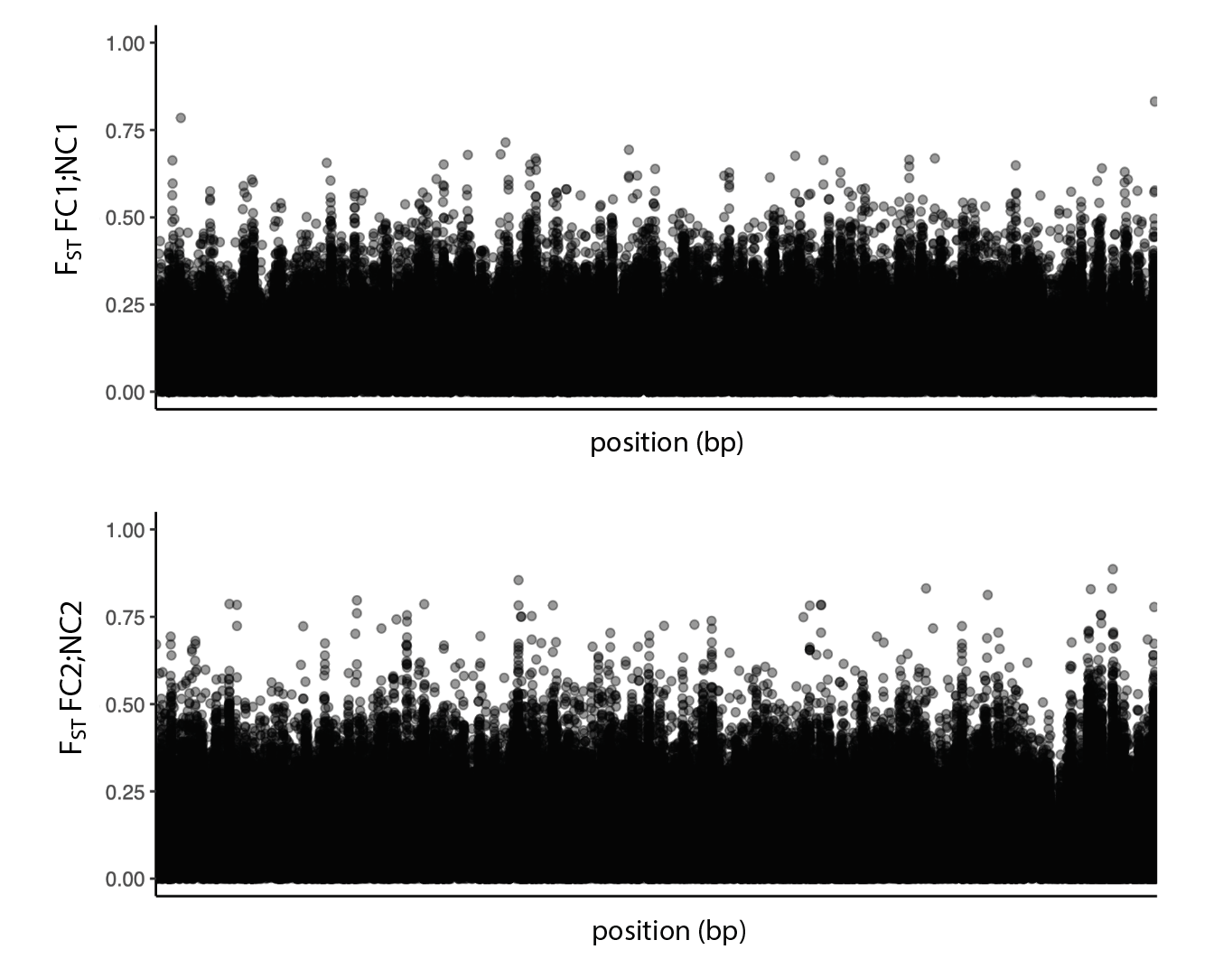

**Figure S2.** Genome-wide F_ST_ between No Care (NC1 and NC2) and Full Care (FC1 and FC2) populations for each replicate block (top: Block 1; bottom: Block 2). Each point represents F_ST_ values for a 500bp sliding window (250bp overlap).

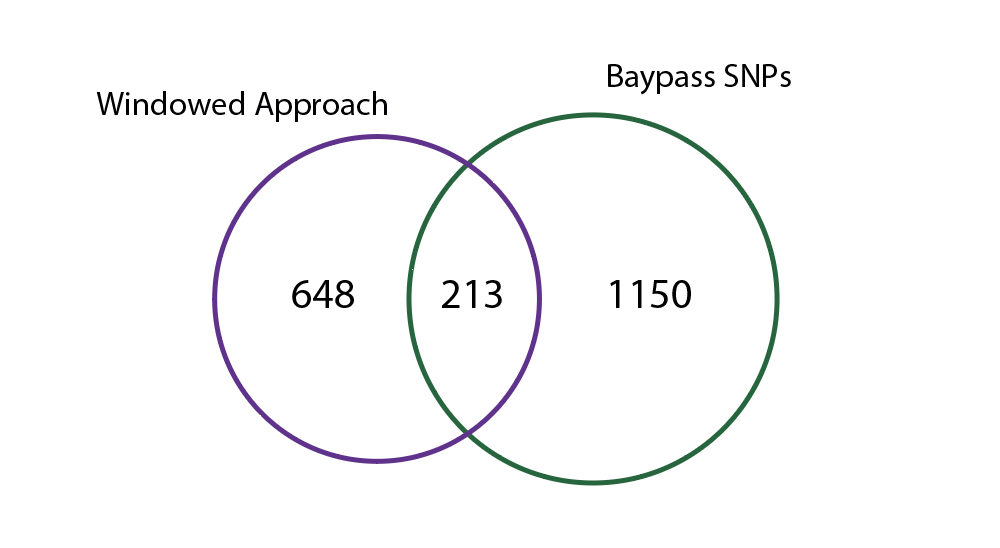

**Figure S3.** Overlap of genes identified between windowed approach (500bp sliding windows using Fishers’ Exact Tests) and using SNP (Baypass auxillary model).

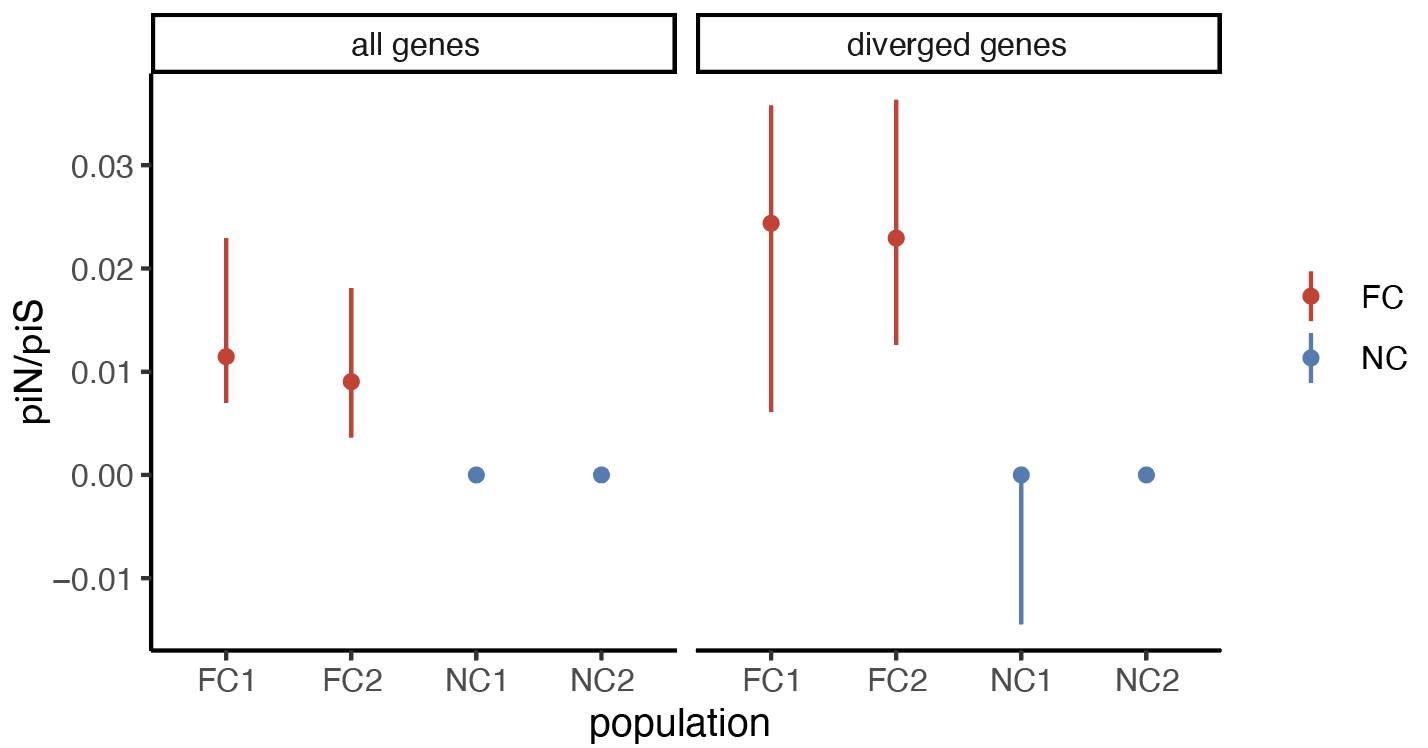

**Figure S4.** Plot of median ratio of non-synonymous to synonymous pi (piN/piS) and boostrapped 95% confidence intervals (CI; error bars) for all genes as well as those that diverged between populations (top 0.5%; n=648). FC = Full Care populations and NC = No Care populations; numbers indicate replicate block.

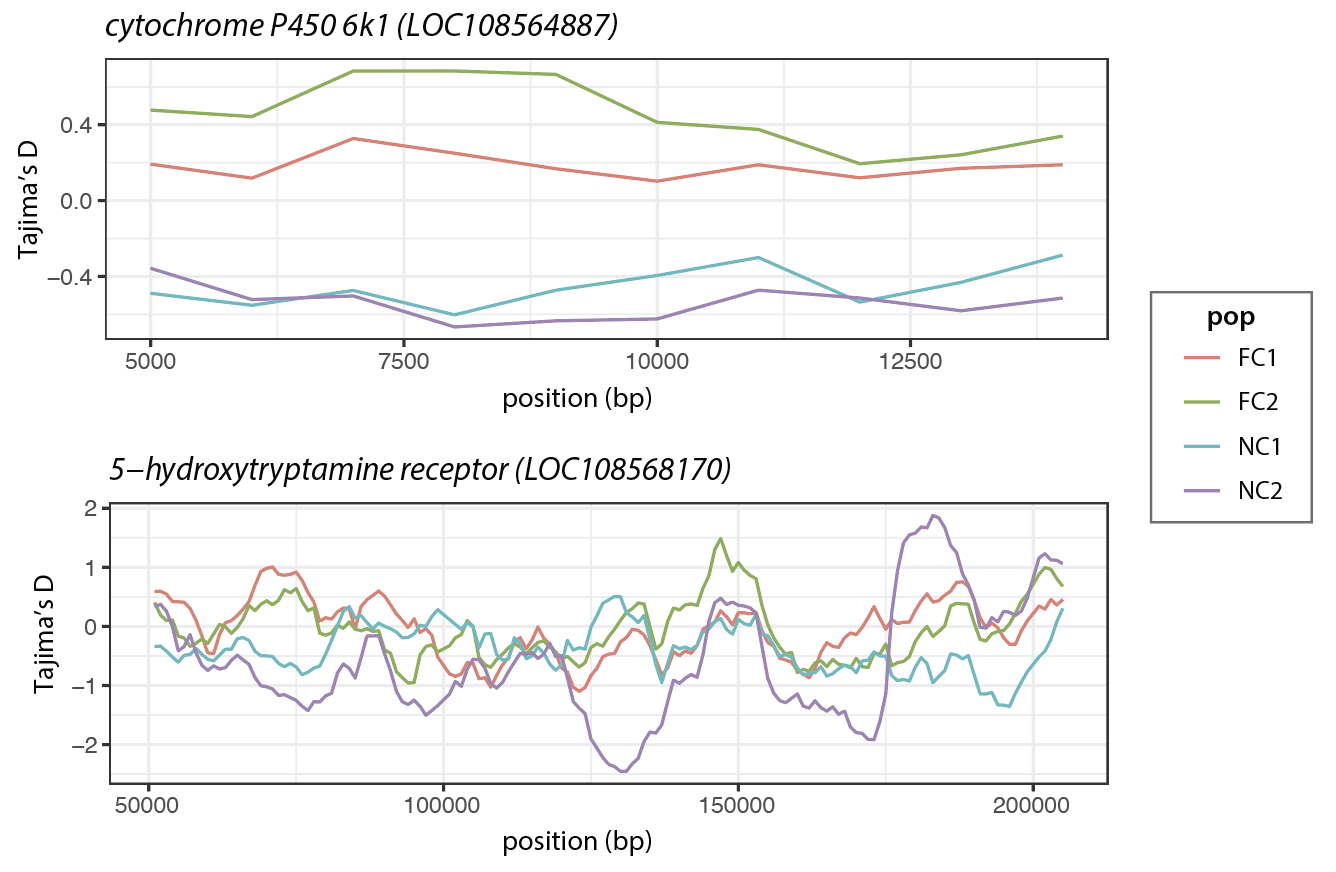

**Figure S5**. Tajima’s D across gene bodies for each population across the gene bodies of (a) cytochrome P450 6k-1 and (b) 5-hydroxytryptamine.
